# Weighted variance component test for the integrative multi-omics analysis of microbiome data

**DOI:** 10.1101/2024.06.14.599073

**Authors:** Angela Zhang, Wodan Ling, Amarise Little, Jessica S. Williams-Nguyen, Jee-Young Moon, Robert D. Burk, Rob Knight, Dong D. Wang, Qibin Qi, Robert C. Kaplan, Ni Zhao, Michael C. Wu

## Abstract

Metabolic dysregulation and alterations have been linked to various diseases and conditions. Innovations in high-throughput technology now allow rapid profiling of the metabolome and metagenome — often the gene content of bacterial populations -– for characterizing metabolism. Due to the small sample sizes and high dimensionality of the data, pathway analysis (wherein the effect of multiple genes or metabolites on an outcome is cumulatively assessed) of metabolomic data is commonly conducted and also represents a standard for metagenomic analysis. However, how to integrate both data types remains unclear. Recognizing that a metabolic pathway can be complementarily characterized by both metagenomics and metabolomics, we propose a weighted variance components framework to test if the joint effect of genes and metabolites in a biological pathway is associated with outcomes. The approach allows analytic p-value calculation, correlation between data types, and optimal weighting. Power simulations show that our approach often outperforms other strategies while maintaining type I error. The approach is illustrated on real data.

## 1 Importance

Advances in sequencing technology have opened up the opportunities for researchers to collect data on multiple aspects of the microbiome. For example, the metagenomics of the microbiome represent can represent metabolic potential whereas its metabolites can represent the metabolic output. Despite these high-throughput innovations, it is still unclear how we can utilize a multiomics approach on these data types to answer our scientific questions of interest, especially in the presence of high-dimensionality. To answer this, we propose a pathway-level testing procedure that uses both the metagenomics and metabolomics in order to assess if a specific microbial pathway is associated with an outcome of diseases, such as disease type. Through both simulations and a real data application, we show that we can achieve better power and discover more significant pathways when we utilize both data types (i.e. metagenomics and metabolomics).

## 2 Introduction

Metabolism represents a central biological component that underlies a wide range of processes in cells and broader organisms. Altered metabolism and metabolic pathways represent a hallmark of multiple diseases and health conditions including diabetes, inflammatory bowel diseases, and certain cancers, among countless others [1, 2, 3]. Consequently, many studies across a range of high-throughput technologies are all focused on understanding the relationship between metabolic processes and a specific outcome of interest. Such studies include everything from genetic associations studies [4] and gene expression profiling studies [5] to metagenomic studies [6] and metabolomic profiling studies [7] However, each of these different data types reflects just one aspect of metabolism. We may be able to achieve greater power if we instead consider the cumulative effect of metabolism characterized through multiple data types.

Due to the small sample sizes and high dimensionality of metabolomic and metagenomic data, pathway analysis (wherein the effect of multiple metabolites on an outcome is cumulatively assessed) of metabolomics data is commonly conducted and also represents the standard for metagenomics analysis. Importantly, metabolic pathways are natural, common units of analysis for both data types. Furthermore, pathway-level analysis reduces the dimensionality of the data (mitigating stringent type I error rate levels). Another advantage to taking a pathway approach as opposed to a gene-by-gene approach is that it better reflects the biological system since cellular and biological processes occur through the concerted activity of multiple features in a pathway. Thus, in terms of biological relevance, an individual feature with a high level of differential abundance may not be as revealing than a group of features with moderate differential abundance.

Several methods have been proposed for pathway-based analyses in metagenomics and metabolomics studies. Simple linear regression models can be used to assess the relationship between the metagenomic pathway (which is typically the total number of sequencing reads mapping to genes within a pathway) and an outcome of interest. Methods for pathway analysis in metabolomic data have largely been adapted from gene expression analysis approaches. One widely used example is over-representation analysis (ORA); in ORA, a list of over-represented pathways is generated from a subset of metabolites (e.g. determined to be statistically significant by some metric) [8]. As an extension of ORA, second and third generations of methods have also been developed for metabolomic pathway analysis. Originally designed for microarray analysis, functional class scoring (FCS) represents the second generation of tools and takes an *a priori* set of metabolites (e.g. all metabolites in a specific pathway) and compares it to an ordered list of metabolites (e.g. all statistically significant metabolites corrected for multiple testing and ranked by log-fold change) to determine whether the *a priori* group of genes are randomly distributed within the list [9]. Network-based analyses make up the third generation and have been shown to be superior to the previous methods since they consider metabolite interactions; topology-based approaches compare known cellular processes and pathways to a network of metabolite to determine if such pathways are altered in a disease condition [10].

We note that the aforementioned methods all utilize the competitive null hypothesis. For a particular pathway, the competitive null hypothesis compares if the genes in the pathway of interest are at most as differentially expressed/abundant as genes/metabolites not in the pathway [11]. Typically, our main objective is to detect metabolic pathways which differ across phenotypes irrespective of how they compare to other pathways, for which the competitive null hypothesis is not suitable[12]. In contrast, a self-contained null hypothesis posits that the metabolites/genes in the selected set *S* is not differentially expressed across phenotypes. Several methods have been developed under the self-contained null hypothesis. Averaging-based approaches such as principal components analysis (PCA)-based methods have been used to select metabolites that capture the most variation in the data set [13, 14]. Although these methods help overcome high-dimensionality in microbiome data, it is difficult to interpret the results from PCA analysis and there is a lack of consensus on how many PC directions need to be selected for analysis. Variance component- and kernel-based approaches have also been proposed for metabolomic pathway analysis. Instead of focusing on each individual metabolites, these methods test if a group of metabolites, such as those found in a specific pathway, are associated with an outcome [15]. However, these approaches are focused on just a single data type. Yet, researchers studying metabolic processes are interested in relating outcomes to metabolic pathways across multiple data sets that collectively represent different aspects of metabolism.

The ability to characterize metabolism using multiple data types offers an opportunity for improving power to identify metabolic relationships on outcomes. As a result, many studies have started collecting both metagenomic and metabolomic data with the objective of relating metabolic pathways to outcomes. In describing integrative metabolomic and metagenomic studies, we can separate the cases where metabolites and metagenomic data are collected from the same specimens or site, such as fecal metabolites and fecal metagenomic samples [16], from the situation in which the metabolites and metagenomic data are collected from different sites, such as when fecal metagenomics and blood metabolites are available[17]. In the former case, many metabolites are of microbial origin such that the metagenome represents the metabolic potential while the metabolome represents the actual metabolic output. These studies can elucidate the host-microbe interactions that play important roles in host metabolism. On the other hand, when data are collected from different sites, then they represent more distinct views of the system. The data are still complementary approaches to characterizing metabolism, but the metabolome-metagenome interactions are expected to be more distal and weaker. Under both cases, the ability to integratively evaluate the effects of data types offers potential for improved statistical power to identify metabolic pathways associated with outcomes; however, the actual analysis of combining both datasets, regardless of the sample site, is challenging.

Several methods have been proposed for an integrative approach to microbiome data analysis [18] . The most popular strategy is to perform a marginal correlation analysis between two biological features. The relationship between these two data sets can be quantified by some pre-specified correlation measure (e.g. Pearson’s correlation, Spearman Rank Correlation) [19]. Although not as popular as marginal correlation analyses, regression-based methods have also been suggested as a potential strategy for elucidating the relationships between two features. This integrative strategy can be framed as a regression model where one feature type, such as the pathway abundance of the metagenomic data, is defined as the response variable and the other feature type, such as the metabolite levels associated with that pathway, can act as the predictor variables. These methods are not limited to only one predictor variable, although sparsity and rank constraints must be employed for multiple response variables [20]. One major limitation to regression-based approach is that they require that one feature type be defined as the predictor and the other as the response variable. This can pose a problem, especially when the underlying biology is not well understood. Furthermore, both marginal correlation-based and regression-based approaches only assess the relationship between feature types. They do not test if an outcome is associated with multiple data sets. For example, these methods are not applicable if we are interested in the relationship between metabolic pathways, represented both by metagenomics and metabolomics data, and disease type.

To address this question, we propose a flexible and computationally efficient regression approach to test association between the joint effect of genes and metabolites in a metabolic pathway and an outcome of interest. In order to account for the high dimensional nature of this problem, we use a weighted variance component (VC) framework and treat each genomic pathway and its corresponding metabolite as a random effect. This setup allows us to achieve reasonable power for representing multiple aspects of metabolism via the metabolic potential (metagenomics) and the metabolic output (metabolomics). Our proposed method is intended for metagenomics data where pathway-level information is available, either from shotgun metagenomics sequencing or inferred through 16S rRNA sequencing. Since the contribution of each data set to the outcome is unknown, we treat our test statistic as a weighted average of the metagenomics score test statistic and metabolomics score test statistic. We perform a grid search on this weighted sum and introduce two potential methods for combination testing for the vector of *p*-values generated from the grid search. Overall, our weighted VC method is a computationally efficient approach and allows for analytic p-value calculation, correlation between data types, and optimal weighting.

The remainder of this manuscript is organized as follows. In the next section, we introduce notation and present our weighted testing procedure by describing the test statistic and the asymptotic null distribution. Then in Section 3, we illustrate the advantages of our proposed approach through simulations and demonstrate the increase in power when considering both representations of metabolism. In Section 4, we show the utility of the weighted VC framework with data from the Hispanic Community Health Study/Study of Latinos (HCHS/SOL) by discerning the subset of metabolic pathways significantly associated with diabetes mellitus (DM). We conclude with a brief discussion in Section 5.

## 3 Weighted variance component test

In this section, we will first introduce notation for the regression model between the outcome of interest and the metagenomics and metabolomics. We will then outline the test statistic from the weighted VC framework and provide two strategies for obtaining the p-value of the test statistic. Finally, we will end the section by providing a kernel-based extension to our proposed method.

### 3.1 Model

Suppose there are *n* subjects that have both metabolomic and metagenomic data for a specific pathway. Let *Y*_*i*_ be a continuous or dichotomous trait of subject *i*. We consider the following regression model:

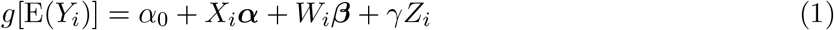

*g*(·) is a link function that is the identity function for continuous outcomes and the logit function for dichotomous outcomes. ***α*** = (*α*_1_, *α*_2_, …, *α*_*p*_) are regression coefficients for covariates, *X*_*i*_ = (*x*_*i*1_, *x*_*i*2_, …, *x*_*ip*_), that we want to adjust out, such as age and sex. *W*_*i*_ = (*w*_*i*1_, *w*_*i*2_, …, *w*_*im*_) is a vector of metabolite levels, and *Z*_*i*_ is the univariate metagenome abundance for a specific pathway. ***β*** = (*β*_1_, *β*_2_, …, *β*_*m*_) and *γ* are the regression coefficients for the *m* metabolites and metagenome abundances respectively. For simplicity, we focus on a single pathway with the understanding that this modeling approach can be applied to each pathway under consideration, in turn.

It is possible for *Z*_*i*_ and *W*_*i*_ to be correlated. For example, if we have a large value *Z*_*i*_, we could expect high levels of metabolites for that particular pathway since a subset of metabolites are synthesized downstream of gene expression. Thus, to decorrelate the two sets of features, we can regress the effect of *Z* from *W* . Specifically, we can rewrite Equation (1) as:

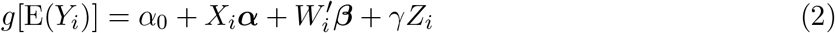

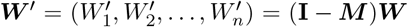 and ***M*** = ***Z***(***Z***^T^***Z***)^−1^***Z***^T^ where ***M*** is a projection matrix onto the column space of ***Z***. Here, 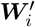 represents the residuals of performing a regression model between each metabolite *W*_*i*_ and the metagenomics value *Z*_*i*_. We are able to achieve better power and obtain asymptotic p-values much more easily by transforming ***W***_*i*_ to be uncorrelated with =*Z*_*i*_.

### 3.2 Weighted variance component test

Evaluating if the metabolome and the metagenome jointly has an effect on the outcome corresponds to testing the null hypothesis: *H*_0_ : *β*_*j*_ = *γ* = 0; *j* = 1, …, *m*. However, this framework assumes independence between the metagenomic and metabolomic datasets and is underpowered for pathways with many metabolites. Instead, suppose that ***β*** is a random variable where each *β*_*j*_ is independent and has mean 0 and variance *ρτ* . Additionally, we suppose that *γ* is a random variable with mean 0 and variance (1 − *ρ*)*τ* . We can see that testing *H*_0_ : *τ* = 0 is equivalent to *H*_0_ : *β*_*j*_ = *γ* = 0. We can use the weighted variance component test for this hypothesis.

Let ***W*** ^*′*^ be a *n* × *k* matrix of the metabolite values that are uncorrelated with *Z*_*i*_ and let ***Z*** be a *n*-length vector of the pathway counts. Furthermore, let 0 ≤ *ρ* ≤ 1. For each *ρ*, we can write the score test statistic as a weighted average of the metabolic potential (metagenome) and the metabolic output (metabolome).

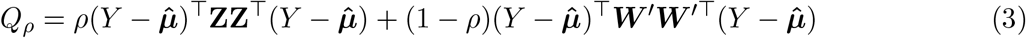

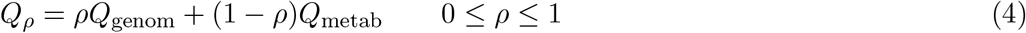

In addition to being computationally advantageous, the score test is preferable over the likelihood ratio test because it only requires fitting the null model; the distribution of the likelihood ratio can be unstable, especially when the estimate is near the boundary. Furthermore, *p*-degrees-of-freedom LR tests can be less powerful when the number of taxa increases and are not even applicable when the number of species exceeds the sample size.

For a fixed value of *ρ*, we can use the usual VC framework. *Q*_*ρ*_ follows a mixture of *χ*^2^ distributions. The p-value for this test statistic can be computed efficiently. Let (*ρ*_1_, …, *ρ*_*j*_) be the eigenvalues of the matrix:

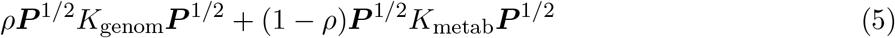

Where 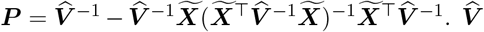 is a diagonal matrix of estimated variance of ***Y*** under the null hypothesis and 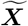 is a *n* × (*p* + 1) matrix [**1, *X***]. Furthermore, *K*_genom_ = **ZZ**^T^ and *K*_metab_ = ***W W*** ^T^. Under this framework:

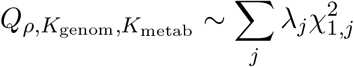

*Qρ*,*K* _genom_,*K*_metab_ is a mixture of independent and identically distributed 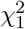 variables. We can calculate the p-value through the Davies method or moment matching method.

Unfortunately, the best value of *ρ* to use is unknown. Therefore, we propose two different strategies to address this barrier. We can calculate a p-value for each value of *ρ* from the grid search by using the previously detailed method. The minimum p-value method (Min p) uses the minimum p-value from this approach as its test statistic. The second method, the Cauchy combination test, employs advantageous features of the Cauchy distribution by using a weighted average of the transformed p-values for its test statistic.

### 3.3 Minimum p-value method

We can see that when *ρ* = 1, the test statistic only involves the metabolomic data and when *ρ* = 0, the test statistic only involves the metagenomic data. In practice, *ρ* is unknown and needs to be estimated for maximal power. To address this, we can perform a grid search; we set a grid of weights, 0 *< ρ*_1_ … *< ρ*_*b*_ *<* 1, to obtain a p-value for each *ρ*_*b*_. In order to select *ρ* that maximizes power, Lee et. al. proposed an optimal test procedure for the generalized SKAT method [21]. Based on this step, we define our test statistic in the following manner:

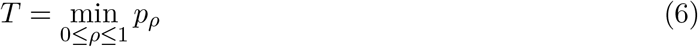

*Q*_metab_ and *Q*_genom_ can be approximated as a mixture of chi-square distributions. In order to obtain the null distribution for our test hypothesis, let 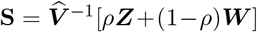 and 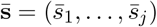 where 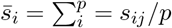. We will define *a*(*ρ*) as:

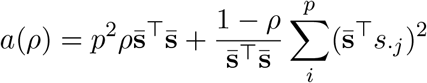

*s*_·*j*_ is the *j*-th column of **S**. For a given *ρ, Q*_*ρ*_ is a mixture of two quadratic forms and is asymptotically equivalent to a mixture of two independent χ^2^ random variables:

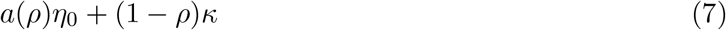

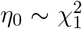 and *κ* approximately follows a mixture of χ^2^. Let *q*_*ρ*_(*T*) be the quantile function for *Q*_*ρ*_. Since *κ* can be approximated with moment matching, we can calculate the *p*-value of *T* :

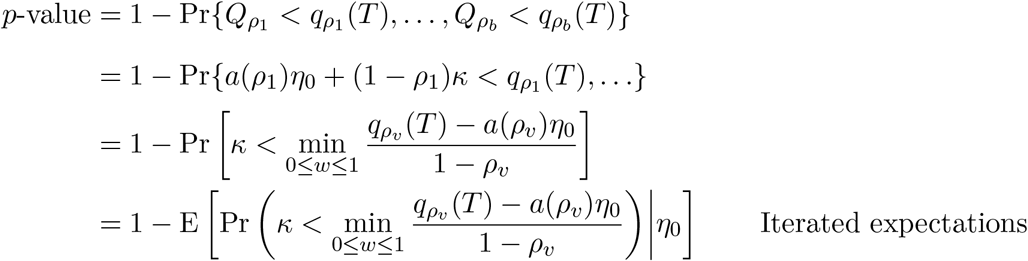

### 3.4 Cauchy combination test

An alternative to the Min-P approach is the Cauchy combination test. P-value combination approaches are primarily used in meta-analyses to aggregate p-values from independent studies that share the same statistical hypothesis. The Fisher method is a popular combination test approach that utilizes the *χ*^2^ distribution to test the null hypothesis of no effect in all studies (citation). We can implement combination testing methods by treating each combination of *ρ* as a separate study.

There have been multiple extensions prescribed for the Fisher method but they often ignore correlation between *p*-values and can be computationally intensive [22, 23, 24]. The Cauchy combination test is a computationally efficient method used for combining individual *p*-values. There are several advantages to this method and it is well-suited to dealing with challenges arising from correlation and high-dimensionality. Under the null hypothesis, *p*-values are uniformly distributed; we can use this fact to transform them into Cauchy-distributed variables. Let (*p*_0_, *p*_0.01_, *p*_0.02_, …, *p*_1_) be a vector of *p*-values from the grid search in the weighted VC test and let *w*_*ρ*_ be the weights associated with each *p*_*ρ*_. The test statistic for the Cauchy combination test is then a weighed sum of Cauchy-transformed *p*-values.

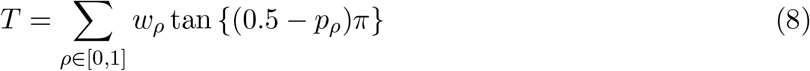

Regardless of the correlation structure of *p*_*ρ*_, *T* has a standard Cauchy distribution under the null. Interestingly, it has been shown that correlation between *p*-values has minimal impact on the tail of the distribution, making this method very powerful for highly-correlated *p*-values. With *w* = Σ_*ρ*_ *w*_*ρ*_, the *p*-value of *T* can be approximated as:

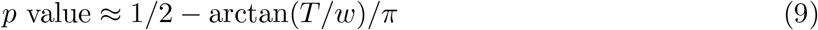

### 3.5 Extension to kernel-based approaches

So far, we have assumed a linear association between the outcome of interest and the metagenome and metabolome for each pathway. Kernel functions are a particularly powerful approach for modeling non-linear and other complex relationships. We can reframe our proposed weighted VC approach in terms of kernel functions:

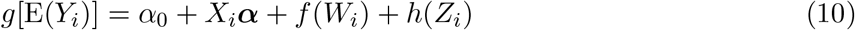

Let *f, h* ∈ *F* a reproducing kernel Hilbert space generated by positive semidefinite kernel functions, *K*(·, ·). Furthermore, let *f* (*W*_*i*_) ∼ *F*_1_(0, *τ*_1_*K*_1_) and *h*(*Z*_*i*_) ∼ *F*_2_(0, *τ*_2_*K*_2_) where *F*_1_ (or *F*_2_) is an arbitrary distribution with mean 0 and variance *τ*_1_*K*_1_ (or *τ*_2_*K*_2_). The use of kernel functions allows flexible modeling of the complex relationship between the outcome and a specific metabolic pathway. At its basis, the kernel function measures the similarity between subjects *i* and *i*^*′*^ based on their data. Let (*ϕ*_1_, …*ϕ*_*b*_) be a basis for *ℱ*. Taking the metabolomics data as an example, we can represent *f* (*W*_*i*_) as a linear combination of the basis functions.

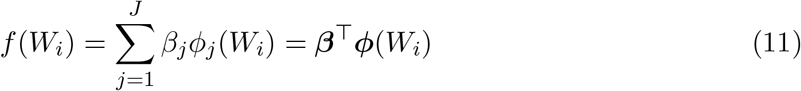

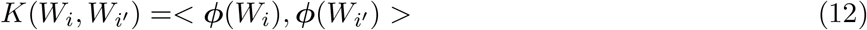

By using the kernel trick, we can implicitly define the basis space by using its inner product, *K*(*W*_*i*_, *W*_*i*_*′*), instead. As a result, more complex models can be specified as long as there is a corresponding kernel function. The test statistic *Q*_*ρ*_ can be rewritten as

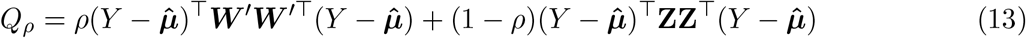

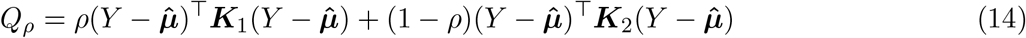

***K***_1_ and ***K***_2_ are kernel functions for the metabolomic and metagenomic data respectively. In the original model, ***W*** ^*′*^***W***^*′*T^ and ***ZZ***^T^ are examples of linear kernels. Although we evaluate the performance of our method with the linear kernel, using a kernel that better captures relationship between the metabolic pathway and the outcome of interest can increase the power of our testing procedure. Pathway data can be represented as a graph where the nodes are the metabolites and the edges are the biological interaction between the metabolites. One such example of this application is the Pathway Integrated Regression-based Kernel Association Test (PaIRKAT), which incorporates prespecified pathway information into its kernel matrix [25].

## 4 Simulations

### 4.1 Simulation settings

We performed several simulations to assess the performance of the weighted VC method. We simulated metagenomic and metabolomic datasets from the empirical cumulative distribution functions of fecal whole-genome shotgun sequencing and capillary electrophoresis time-of-flight mass spectrometry (CE-TOFMS)-based metabolomics collected from 616 participants at the National Cancer Center Hospital in Tokyo, Japan. Metabolites were grouped based on the corresponding Kyoto Encyclopedia of Genes and Genomes (KEGG) pathway given by the metagenomics data. In order to gauge the performance of the proposed method over pathways of varying size, we simulated data from two pathways: glycerophospholipid metabolism (map00564) which consisted of 6 metabolites and ABC transporters (map02010) which contained 37 metabolites. We tested the performance of our method on both continuous and dichotomous outcomes of interest and used a grid of ***ρ*** = (0, 0.01, 0.02, …, 0.99, 1.00) for our weights.

### 4.2 Type I Error

To evaluate type I error for continuous and dichotomous outcomes, we simulated 5000 datasets each for *n* = (50, 100, 200, 300, 400). In the following models below, we set ***α*** = (1, 0.5, 0.5) and **X** as a matrix of covariates that we wanted to adjust out. We evaluated the performance of our proposed method on both continuous and dichotomous outcomes (*Y*_*i*_). In order to test the performance of adjusting for both continuous and dichotomous covariates, *X*_1_ was generated from a standard normal distribution and *X*_2_ was generated from Bernoulli(0.5). The error terms, *ϵ*_*i*_, were generated from independent standard normal distributions. We defined type I error as the proportion of *p*-values less than ***α***_p-value_ = (0.01, 0.05). Under the null model, we assumed the following models for continuous and dichotomous outcomes respectively:

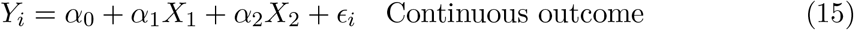

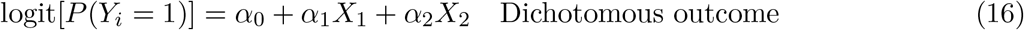

Type I performance of the weighted VC method for these simulations are shown in Table 1. Overall, the weighted VC method agrees well with ***α***_p-value_ = (0.01, 0.05) for both pathways and types of outcomes. Our proposed method tends to be slightly conservative for small sample sizes in the continuous case, although this deviation is small. We note that the weighted VC test has slightly inflated Type I error in the dichotomous outcome scenario when the sample size is small, but this modest deviation decreases as sample size increases. Type 1 error results in the dichotomous case are included in the supplemental section (see Table 4 and 5).

**Table 1:** Type I error rates for the *α* = (0.01, 0.05) and *n* = (50, 100, 200, 300, 400) for map02010 in the continuous outcome case.

| **n** | **Min-P** |  | **Cauchy** |  |
| --- | --- | --- | --- | --- |
| | $\alpha = \mathbf{0.05}$ | $\alpha = \mathbf{0.01}$ | $\alpha = \mathbf{0.05}$ | $\alpha = \mathbf{0.01}$ |
| **50** | 0.046 | 0.0078 | 0.0496 | 0.0082 |
| **100** | 0.046 | 0.088 | 0.0452 | 0.0096 |
| **200** | 0.048 | 0.0094 | 0.0502 | 0.0092 |
| **300** | 0.0512 | 0.0102 | 0.0544 | 0.011 |
| **400** | 0.0466 | 0.0074 | 0.0464 | 0.0072 |

**Table 2:**
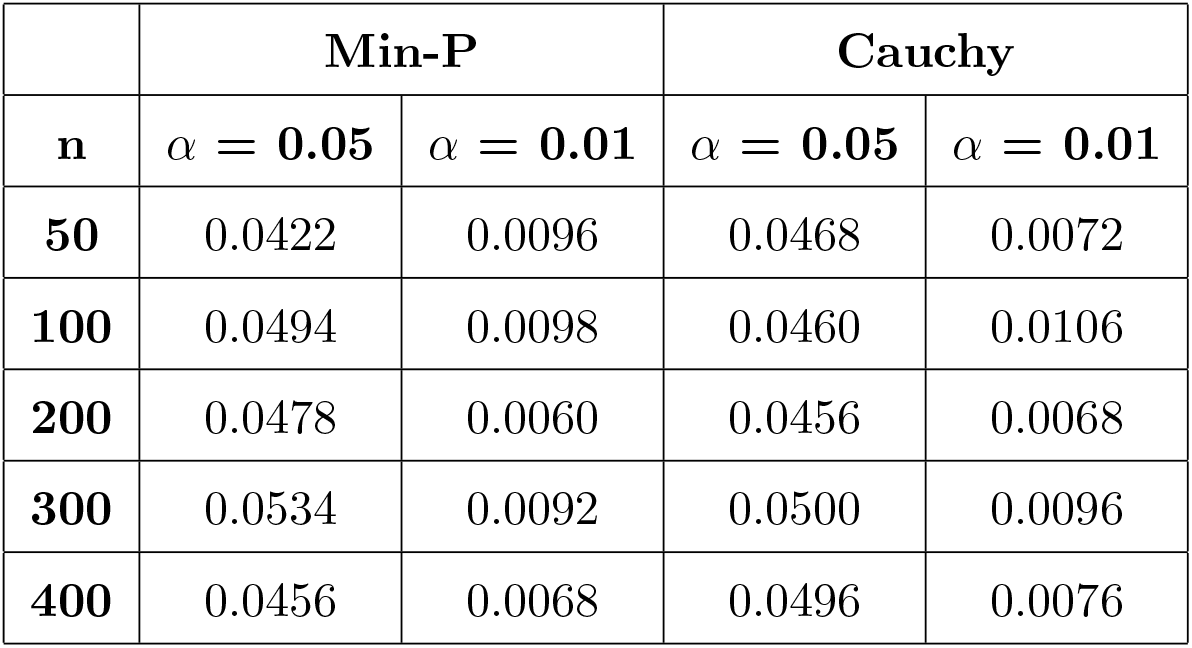
Type I error rates for the *α* = (0.01, 0.05) and *n* = (50, 100, 200, 300, 400) for map00564 in the continuous outcome case.

|  | Min-P |  | Cauchy |  |
| --- | --- | --- | --- | --- |
| **n** | $\alpha = 0.05$ | $\alpha = 0.01$ | $\alpha = 0.05$ | $\alpha = 0.01$ |
| **50** | 0.0422 | 0.0096 | 0.0468 | 0.0072 |
| **100** | 0.0494 | 0.0098 | 0.0460 | 0.0106 |
| **200** | 0.0478 | 0.0060 | 0.0456 | 0.0068 |
| **300** | 0.0534 | 0.0092 | 0.0500 | 0.0096 |
| **400** | 0.0456 | 0.0068 | 0.0496 | 0.0076 |

### 4.3 Power

To evaluate the power of our approach, we conducted simulations under the alternative hypothesis. Specifically, we assumed the model:

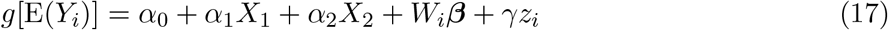

We set ***α***_Power_ = (1, 0.3, 0.3). For the first scenario, we assessed the performance of the metabolic potential while keeping the effects of the metabolic output constant. We set a non-zero constant value for a random selection of ∼ 20% of the ***β*** while the remaining ∼ 80% of the coefficients were set to 0. We first tested the performance of the weighted VC method by varying the effects of *γ*. In the second scenario, we evaluated the performance of the method by keeping the metabolic potential constant while varying the effects of the metabolic output. We set a constant value for *γ* based on the first scenario and calculated the power of the method over a range of different constant values for ***β***. Although we only show the continuous outcome case for both scenarios, we observed similar results in the dichotomous outcome case and are shown in the Supplemental Section (See Figure 3 and 4). We compared the power of our proposed weighted VC test using both integrated methods (Min-p and Cauchy) to methods that utilized only one dataset. For the metagenomics data, each pathway is represented by one value, which is the combination of all the gene expression activity for that specific pathway. Because each pathway contains only one value for each sample, we assumed a simple linear regression model in the continuous scenario:

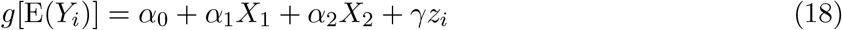

For the metabolomics data, we assumed the association between the metabolites and outcome as:

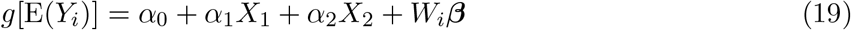

In the continuous scenario, both *g*[E(*Y*_*i*_)] in Equation 18 and Equation 19 are the identity link function. In contrast to the metagenomic data, there can be a modest to large number of metabolites in a pathway. To accommodate possible high-dimensionality of the metabolites, we used a standard VC test for the metabolomics-only analysis. We assumed that each *β*_*j*_ followed an arbitrary distribution with mean 0 and variance *τ* . Our test statistic for evaluating the null hypothesis, *τ* = 0, was:

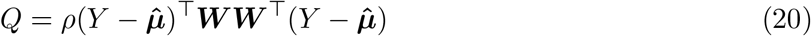

*Q* asymptotically follows a mixture of χ^2^ distributions. Specifically 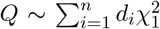 where *d*_*i*_ are the eigenvalues of **Σ**^1*/*2^***P***_*n*_**WW**^T^***P***_*n*_**Σ**^1*/*2^ with **Σ** = cov(***Y***). The Davies method, which involves the numerical inversion of the characteristic function, can be used to obtain a *p*-value for the test statistic. In short, we can view our procedure for assessing the power of a metabolomics-only analysis as a simplified version of the proposed weighted VC test.

We present the results of Scenario 1 in Figure 1. For map00564, the metabolomics-only analysis only outperformed the integrated methods when there was no metagenomics effect, i.e. *γ* = 0. While the power of the metabolomics-only analysis decreased as the effect size of *γ* increased, the performance of the metagenomics-only analysis rapidly improved. For *γ >* 0.1, the integrated methods consistently outperformed the performance of the metagenomics-only and metabolomics-only analysis for all sample sizes. With regards to the integrated methods, the Cauchy and Min-p approach had similar performance unless the sample size was small and the effect size of *γ* was large or small. Recall that the Min-p method relies on one *p*−value while the Cauchy method is an average of Cauchy-transformed *p*−values. As a result, the Min-p method will be especially advantageous when the contribution of each dataset is unequal, i.e. one data set has a dominating effect on the outcome. Similar results from the map00564 pathway were also observed in the map02010 pathway.

**Figure 1.**
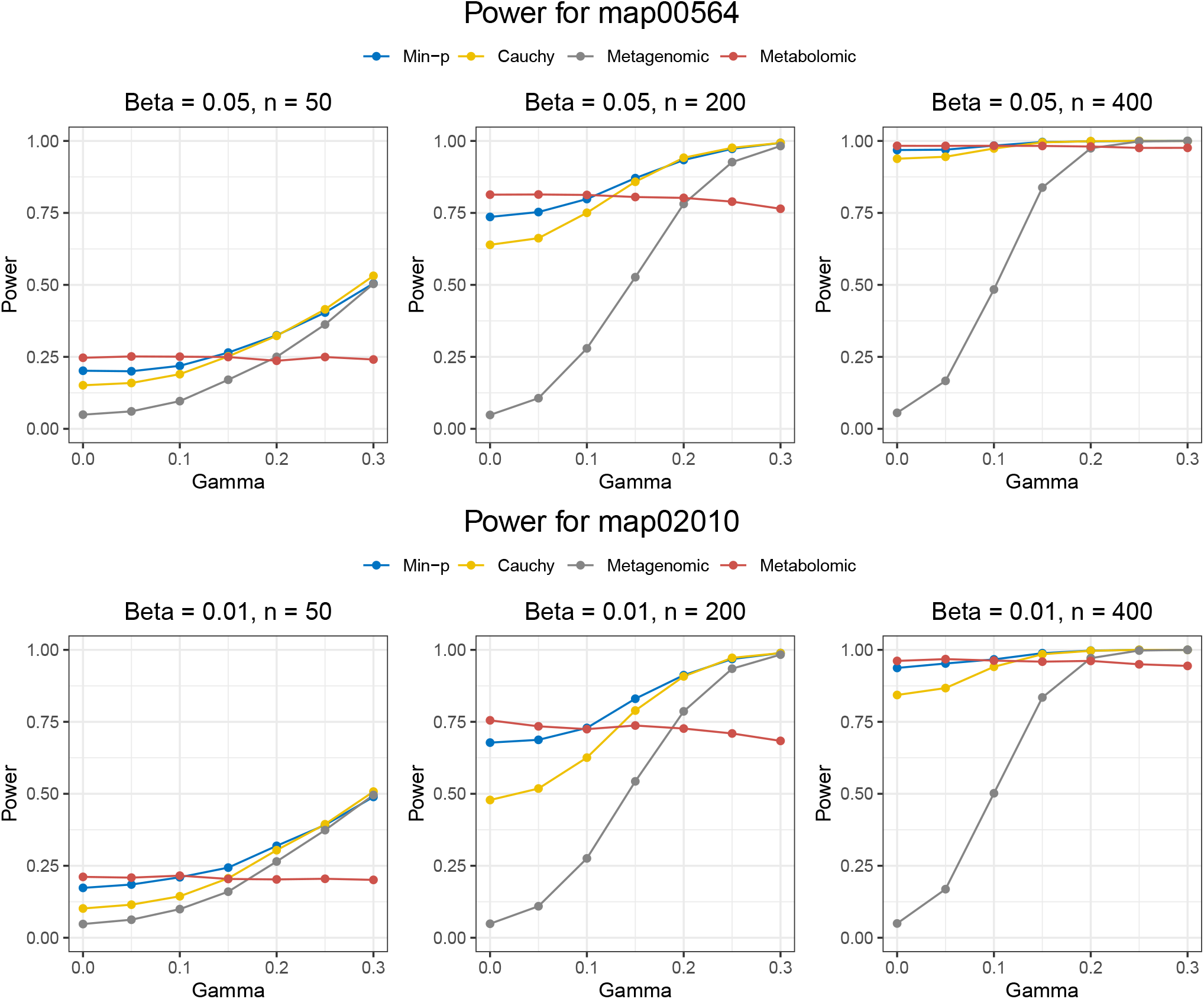
Simulation results for evaluating power in Scenario 1. The effects of the metabolomics input is kept constant as we evaluate the impact of increasing the effects of the metagenomics input for *n* = (50, 200, 400) and for pathways map00564 and map02010. Power is defined as the proportion of *p*−values ≤ 0.05

For the second scenario, we chose a value of *γ* that had moderate power in Scenario 1 and then varied the effects of ***β***. The results of Scenario 2 are outlined in Figure 2. Although the performances of the integrated methods increased as ***β*** increased, the metabolomics-only analysis had the greatest power at *n* = 50 since the contribution between the metabolomics data and metagenomics data becomes unequal; i.e. the metabolomics effect dominates the metagenomics effect. We note that the increase in the metabolomics-only performance is minimal and requires prior knowledge that the metabolites are more strongly associated with the outcome than the metagenes. For an agnostic approach, it is still preferable to use either of the integrated methods. For *n* = (200, 400), both integrated methods outperformed the other two methods across all values of ***β*** for both pathways.

**Figure 2.**
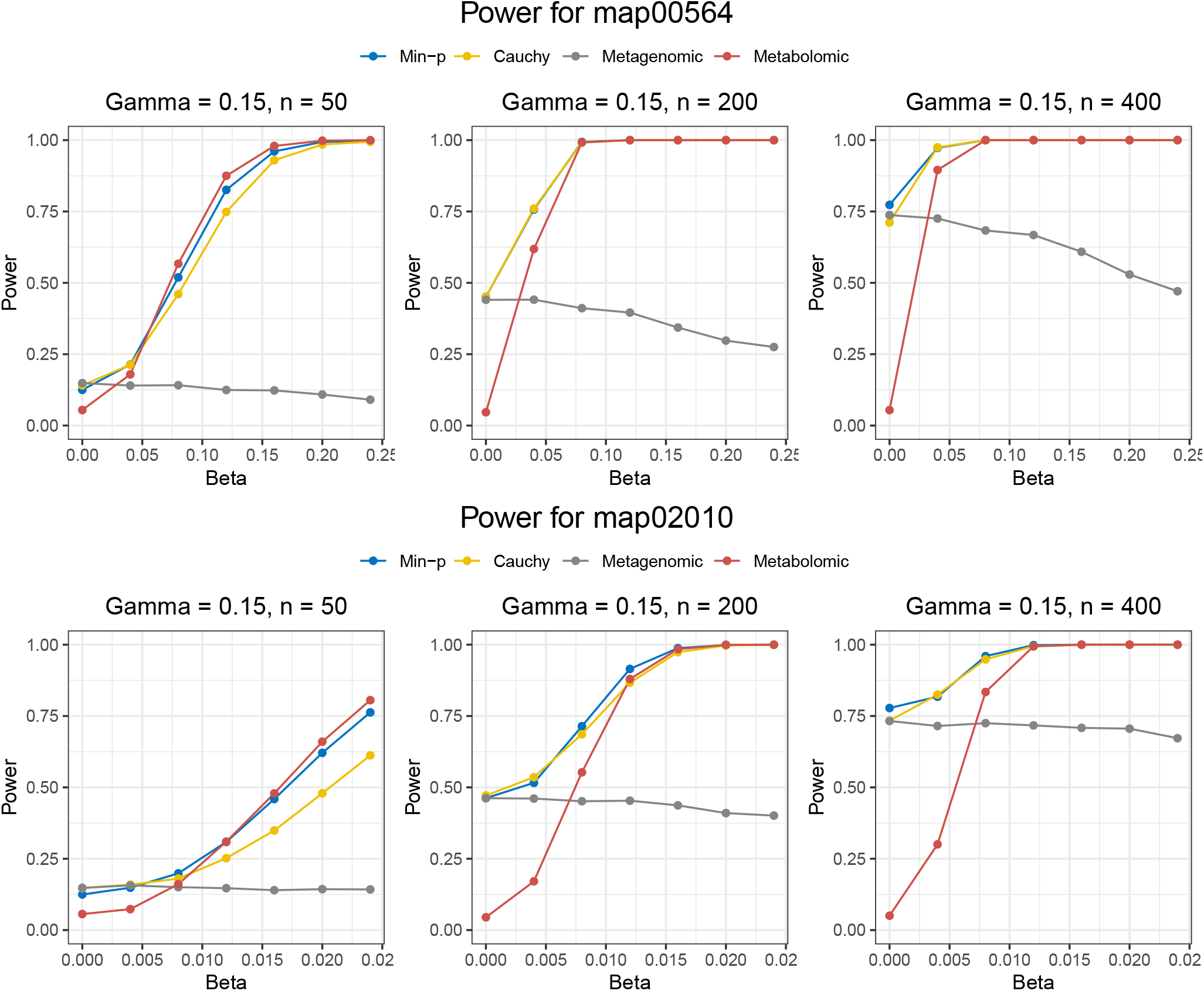
Simulation results for evaluating power in Scenario 2. The effects of the metabolomics input is kept constant as we evaluate the impact of increasing the effects of the metagenomics input for *n* = (50, 200, 400) and for pathways map00564 and map02010. Power is defined as the proportion of *p*−values ≤ 0.05

## 5 Application to the HCHS/SOL Study

The Hispanic Community Health Study/Study of Latinos (HCHS/SOL) is a multicenter epidemiologic study aimed at assessing the risks and determinants of health in Hispanic/Latino populations across the United States. Approximately 16,000 participants across Miami, San Diego, Chicago and the Bronx area of New York are involved in the study. Cardiovascular disease (CVD) is particularly prevalent in Hispanics/Latinos in the United States and is the leading cause of mortality in this population [26]. Several key risk factors for CVD have been identified and include hypertension, high cholesterol levels, diabetes and obesity [27]. Baseline findings of the HCHS/SOL study has shown that a large majority of this population, 71% in women and 80% in men, have at least one major risk factor for CVD [28].

Our work is motivated by a microbiome sub-study of HCHS/SOL. Recent developments in microbiome research have led to increased interest in understanding the crosstalk between the gut microbiome and disease risk. The Centers for Disease Control and Prevention estimate that more than 30 million people in the United States have diabetes mellitus (DM), a major risk factor of CVD. Recent studies have shown the functional role of the gut microbiome in the pathophysiology of T2D. In particular, T2D is characterized by increased permeability of the gut. Reduced alpha diversity, as seen in patients with T2D, can lead to weaker tight junctions that result in increased insulin resistance and chronic inflammation [29]. Given that pathways disrupted by T2D can be contributed by both the host and microbiome, we applied the weighted VC method to these data sets to determine the subset of pathways significantly associated with T2D. Due to its status as a CVD risk factor, identifying significant pathways in T2D not only increases our understanding of this chronic disease but also helps aid in the development of comprehensive and effective preventive measures for CVD.

In our analysis, we used KEGG-based pathways in order to connect each metabolite to its metagenomic pathway. To encourage normality, we log-transformed both datasets. We compared our method with a metagenomics-only and metabolomics-only analysis described earlier in the Simulations section. After data quality control, we had whole genome shotgun sequencing data and targeted serum metabolomic data for 46 pathways from 621 individuals. Of these 621 individuals, 137 had normal glucose regulation, 290 had impaired glucose regulation, and 194 were diagnosed with diabetes. We applied our proposed method on each of the 46 pathways and adjusted our *p*-values using the false discovery rate (FDR) correction. Significant pathways were defined as having FDR ≤ 0.05.

The results of our real-data analysis are shown in Table 3. Cumulatively, this analysis identified many significant pathways associated with T2D. Quality of life is often broadly affected by T2D and there is substantive literature connecting T2D to renal function, coronary arterial disease, retinopathy, neuropathy, depression and dementia among many other conditions [30]. Since diabetes is a condition that affects so many biological processes, it is reasonable to see many pathways to be significant with diabetes status.

**Table 3:** Unique and overlapping significant pathways associated with diabetes mellitus (DM) from the HCHS/SOL Study

|  | Min p-val | Cauchy | Metagenomics | Metabolomics |
| --- | --- | --- | --- | --- |
| Min p | 29 | - | - | - |
| Cauchy | 21 | 21 | - | - |
| Metagenomics | 0 | 0 | 0 | - |
| Metabolomics | 23 | 20 | 0 | 24 |

The Min-p method was able to detect more diabetes-relevant pathways compared to the other three methods. Additionally, as seen in Table 1, there were many overlapping pathways that were identified in Min-p method, Cauchy method and the metabolomics-only analysis; no significant pathways were detected in the metagenomics-only analysis. Based on this result and the large number of overlapping pathways between the Min-p method and metabolomics-only analysis, the differences in metabolic pathways between disease types seem to be driven primarily by the metabolites. However, it is still advantageous to consider both data sets since the Min-p method was able to detect six unique pathways compared to the metabolomics-only analysis. Overall, our analysis has identified significant novel pathways that can be investigated in future studies to elucidate the mechanisms behind both T2D and CVD.

## 6 Discussion

We have proposed a weighted variance component framework for integrating metagenomics and metabolomics data when assessing the association between a pathway and an outcome of interest. In understanding the relationship between the host and the microbiome, metabolism can be represented in two parts: the metabolic potential (metagenomics) from the microbiome and the metabolic output (metabolomics) from the host. By employing a grid search strategy, we are able to agnostically evaluate the weighted average effect of both data sets. Compared to methods that only consider one data, we show through simulations that we actually increase power while also controlling for Type I error if we consider both the metagenome and the metabolome in our analysis. Lastly, we applied our proposed approach to the HCHS/SOL study and identified several unique pathways significantly associated with T2D.

The synthesis of some metabolites from gene expression may suggest the redundancy of using two datasets in our proposed method. However, we present several theories that may explain why we obtain higher power when we consider both data sets. First of all, metabolomic data can often be noisy. In addition to measurement error from the equipment, levels of metabolites are highly variable on the time the sample was collected. A recent study examined the within-individual variability and between-individual variability in 385 metabolites from 60 individuals from baseline and one year into the Shanghai Physical Activity Study. Within-individual variability captured the majority of variation explained for 64% of the observed metabolites [31]. Furthermore, metagenomic data can capture shorter and more volatile metabolites that standard mass spectrometry procedures fail to detect. Lastly, in the context of our real data analysis, we had host metabolites and microbial genes. We can represent both host and microbial mechanisms as a potential application of our proposed approach. Using a weighted VC framework to capture these two aspects of metabolism, we can better assess the association between an outcome and a metabolic pathway. As an extension, our method can be used to evaluate the cumulative effect of any two data sets (e.g. transcriptomics, proteomics, epigenomics, etc.).

We introduce two methods for obtaining *p*-values from our testing procedure: the minimum p-value method and the Cauchy combination test. The former method relies on the result that for each *ρ*, the test statistic *Q*_*ρ*_ can be approximated as the sum of two independent chi-square distributions. Despite this assumption, the Min-p method has been shown to have good performance in small sample sizes. The Cauchy combination test, on the other hand, is based on the result that the weighted sum of Cauchy-transformed *p*-values are still Cauchy under arbitrary correlation structures. Both the minimum p-value method and Cauchy combination test performed well in simulation studies, though we note the former method was able to detect more significant pathways in the real data analysis. Although both methods are suitable in the presence of correlated *p*-values, the main difference between the minimum p-value method and the Cauchy combination test is the representation of the final test statistic as a minimum or as an average, respectively. For example, the minimum p-value method is advantageous if there is a large difference between the minimum and the rest of the elements in the *p*-value vector. However, if there is low variability within the vector, the usage of the Cauchy combination test may lead to increased power. Additionally, we note that the Cauchy combination test is computationally faster and simpler to implement.

Through our simulation studies and real data analysis, we demonstrate that the integration of metabolic potential and metabolic output leads to increased power and the increased detection of significant pathways, respectively. As technology evolves, we now have the opportunity to collect multiple data sets to answer our scientific questions of interest. So far we have applied our method in the context of host and microbial contributions; our method can also be applied to host-only or microbiome-only data sets. Additionally, in our real-data example, we applied a *de novo* approach determining significant pathways, i.e. we used the weighted VC test on all available pathways. Our method may be powerful on a subset of more carefully selected pathways, like those with metabolites known to be relevant to bacterial production or function.

## 7 Supplemental Information

In this section, we show type 1 error and power simulation results for the weighted VC method when we have a dichotomous outcome. This section is adapted from the continuous example in the main text. The mean-generating model for type 1 error is:

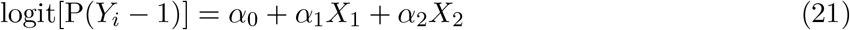

Type 1 error for the dichotomous case are included in Table 4 and 5.

**Table 4:** Type I error rates for the *α* = (0.01, 0.05) and *n* = (50, 100, 200, 300, 400) for map02010 in the dichotomous case.

|  | **Min-P** |  | **Cauchy** |  |
| --- | --- | --- | --- | --- |
| **n** | $\alpha = \mathbf{0.05}$ | $\alpha = \mathbf{0.01}$ | $\alpha = \mathbf{0.05}$ | $\alpha = \mathbf{0.01}$ |
| **50** | 0.0500 | 0.0090 | 0.0500 | 0.0092 |
| **100** | 0.0400 | 0.0108 | 0.0400 | 0.0104 |
| **200** | 0.0200 | 0.0104 | 0.0400 | 0.0096 |
| **300** | 0.070 | 0.0118 | 0.0600 | 0.0100 |
| **400** | 0.0400 | 0.0116 | 0.0500 | 0.0100 |

**Table 5:** Type I error rates for the *α* = (0.01, 0.05) and *n* = (50, 100, 200, 300, 400) for map00564 in the dichotomous case.

|  | **Min-P** |  | **Cauchy** |  |
| --- | --- | --- | --- | --- |
| **n** | $\alpha = \mathbf{0.05}$ | $\alpha = \mathbf{0.01}$ | $\alpha = \mathbf{0.05}$ | $\alpha = \mathbf{0.01}$ |
| **50** | 0.0800 | 0.0112 | 0.0700 | 0.0108 |
| **100** | 0.0600 | 0.0104 | 0.0600 | 0.0108 |
| **200** | 0.0500 | 0.0094 | 0.0400 | 0.0090 |
| **300** | 0.0200 | 0.0092 | 0.0600 | 0.0084 |
| **400** | 0.0600 | 0.0108 | 0.0600 | 0.0116 |

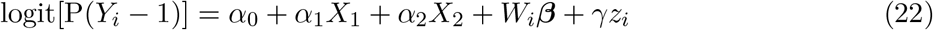

For the first scenario, we assessed the performance of the metabolic potential while keeping the effects of the metabolic output constant. We set a constant value for ≈ 20% of ***β*** while the remaining 80% of coefficients were set to 0. We first tested the performance of the weighted VC method by varying the effects of *γ*. In the second scenario, we evaluated the performance of the method by keeping the metabolic potential constant and varying the effects of the metabolic output. We set a constant value for *γ* based on the first scenario and calculated the power of the method over a range of different constant ***β*** values. We compared the power of the weighted VC test, using both integrated methods (Min-p and Cauchy) to methods that just used one dataset. Since the metagenomics data for each pathway contains just value for each sample, we used a simple logistic regression model for the metagenomic data in the continuous scenario:

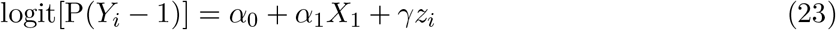

We modeled the association between the metabolites and outcome as:

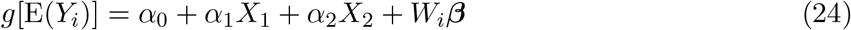

In contrast of the metagenomics-only analysis, we used a variance component test for the metabolomics-only analysis since there can be a modest to large number of metabolites in a pathway. We assumed that each *β*_*j*_ followed an arbitrary distribution with mean 0 and variance *τ* . Our test statistic for evaluating the null hypothesis: *τ* = 0 was:

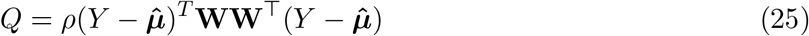

*Q* asymptotically follows a mixture of *χ*^2^ distributions. Specifically 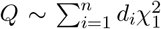 where *d*_*i*_ are the eigenvalues of **Σ**^1*/*2^***P***_*n*_**WW**^T^***P***_*n*_**Σ**^1*/*2^ with **Σ** = cov(***Y***). The Davies method, which involves the numerical inversion of the characteristic function, can be used to obtain a *p*-value for the test statistic. In short, our procedure for assessing the power of a metabolomics-only analysis is a simplified version of the proposed weighted VC test.

As shown in Figure 3, our results from Scenario 1 are largely the same as the continuous outcome example. For map00564, the metabolomics-only analysis only outperformed the integrated methods when there was no metagenomics effect, i.e. *γ* = 0. However, for *γ >* 0.2 the integrated methods consistently outperformed the performance of the metagenomics-only and metabolomics-only analysis for all sample sizes. With regards to the integrated methods, the Cauchy and Min-p approach had similar performance unless the sample size was small and the effect size of *γ* was large or small. Recall that the Min-p method relies on one *p*−value while the Cauchy method is an average of *p*−values. Similar results were also observed in the map02010 pathway.

**Figure 3.**
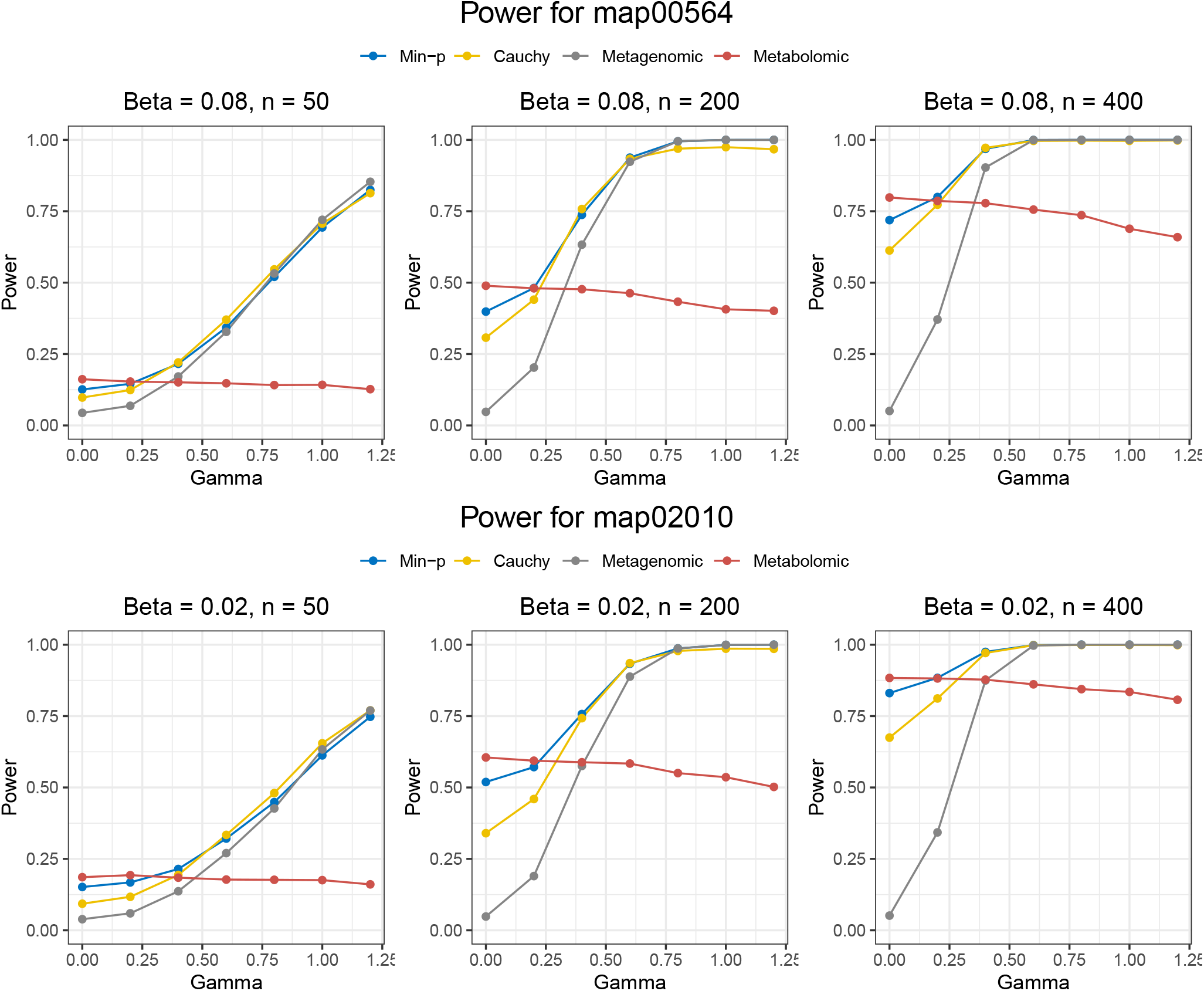
Simulation results for evaluating power in Scenario 1 with a dichotomous outcome. The effects of the metabolomics input is kept constant as we evaluate the impact of increasing the effects of the metagenomics input for *n* = (50, 200, 400) and for pathways map00564 and map02010. Power is defined as the proportion of *p*−values ≤ 0.05

For the second scenario, we chose a value of *γ* that had moderate power in Scenario 1 and then varied the effects of ***β***. The results of Scenario 2 are outlined in Figure 4. Although the performances of the integrated methods increase as ***β*** increases, the metabolomics-only analysis has the largest power at when ***β*** is large since the contribution between the metabolomics data and metagenomics data is unequal. We note that the increase in performance in the metabolomics-only approach is minimal and requires prior knowledge that the metabolites have a greater effect on the outcome than the metagenes. For an agnostic approach, it is still preferable to use either of the integrated methods.

**Figure 4.**
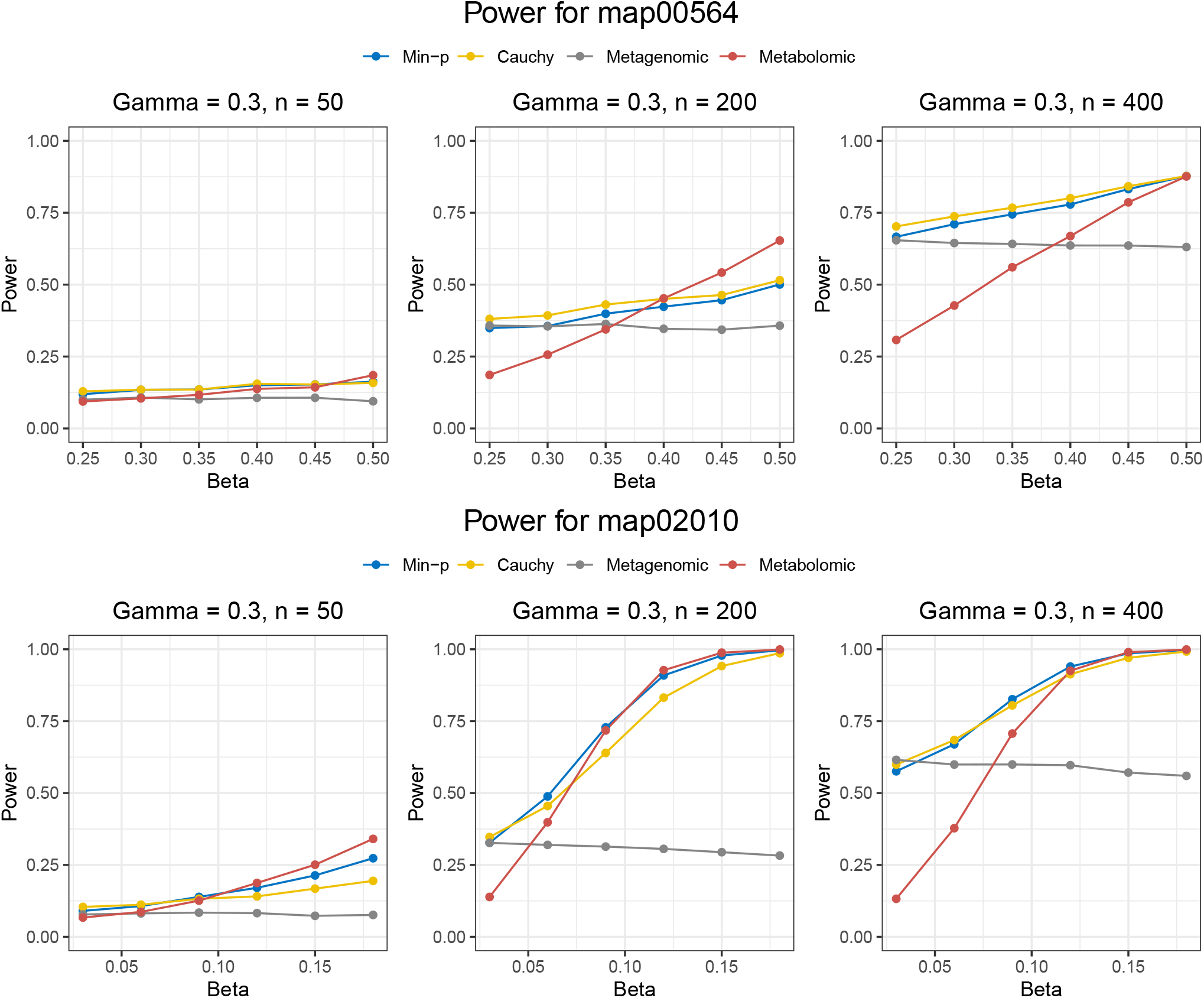
Simulation results for evaluating power in Scenario 1 with a dichotomous outcome. The effects of the metagenomics input is kept constant as we evaluate the impact of increasing the effects of the metabolomics input for *n* = (50, 200, 400) and for pathways map00564 and map02010. Power is defined as the proportion of *p*−values ≤ 0.05

## References

[1] R. R. Kalyani and J. M. Egan, “Diabetes and altered glucose metabolism with aging,” Endocrinology & Metabolism Clinics of North America, vol. 42, pp. 333–347, Jun 2013.

[2] S. Ardizzone, S. Bollani, P. Bettica, M. Bevilacqua, P. Molteni, and G. Bianchi Porro, “Altered bone metabolism in inflammatory bowel disease: there is a difference between Crohn’s disease and ulcerative colitis,” Journal of Internal Medicine, vol. 247, pp. 63–70, Jan 2000.

[3] R. J. DeBerardinis and N. S. Chandel, “Fundamentals of cancer metabolism,” Science Advances, vol. 2, p. e1600200, 05 2016.

[4] J. C. Pardo, V. Ruiz de Porras, J. Gil, A. Font, M. Puig-Domingo, and M. Jordà, “Lipid metabolism and epigenetics crosstalk in prostate cancer,” Nutrients, vol. 14, p. 851, Feb. 2022.

[5] J. Makoukji, N. J. Makhoul, M. Khalil, S. El-Sitt, E. S. Aldin, M. Jabbour, F. Boulos, E. Gadaleta, A. Sangaralingam, C. Chelala, R. M. Boustany, and A. Tfayli, “Gene expression profiling of breast cancer in Lebanese women,” Sci Rep, vol. 6, p. 36639, Nov 2016.

[6] J. Qin, Y. Li, Z. Cai, S. Li, J. Zhu, F. Zhang, S. Liang, W. Zhang, Y. Guan, D. Shen, Y. Peng, D. Zhang, Z. Jie, W. Wu, Y. Qin, W. Xue, J. Li, L. Han, D. Lu, P. Wu, Y. Dai, X. Sun, Z. Li, A. Tang, S. Zhong, X. Li, W. Chen, R. Xu, M. Wang, Q. Feng, M. Gong, J. Yu, Y. Zhang, M. Zhang, T. Hansen, G. Sanchez, J. Raes, G. Falony, S. Okuda, M. Almeida, E. LeChatelier, P. Renault, N. Pons, J. M. Batto, Z. Zhang, H. Chen, R. Yang, W. Zheng, S. Li, H. Yang, J. Wang, S. D. Ehrlich, R. Nielsen, O. Pedersen, K. Kristiansen, and J. Wang, “A metagenome-wide association study of gut microbiota in type 2 diabetes,” Nature, vol. 490, pp. 55–60, Oct 2012.

[7] W. G. Hunter, J. P. Kelly, R. W. McGarrah, W. E. Kraus, and S. H. Shah, “Metabolic Dysfunction in Heart Failure: Diagnostic, Prognostic, and Pathophysiologic Insights From Metabolomic Profiling,” Curr Heart Fail Rep, vol. 13, pp. 119–131, Jun 2016.

[8] J. J. Goeman and P. Bühlmann, “Analyzing gene expression data in terms of gene sets: methodological issues,” Bioinformatics, vol. 23, pp. 980–987, Apr 2007.

[9] A. Subramanian, P. Tamayo, V. K. Mootha, S. Mukherjee, B. L. Ebert, M. A. Gillette, A. Paulovich, S. L. Pomeroy, T. R. Golub, E. S. Lander, and J. P. Mesirov, “Gene set enrichment analysis: a knowledge-based approach for interpreting genome-wide expression profiles,” PNAS, vol. 102, pp. 15545–15550, Oct 2005.

[10] C. Mitrea, Z. Taghavi, B. Bokanizad, S. Hanoudi, R. Tagett, M. Donato, C. Voichiţa, and S. Drăghici, “Methods and approaches in the topology-based analysis of biological pathways,” Front. Physiol., vol. 4, p. 278, Oct. 2013.

[11] M. C. Wu and X. Lin, “Prior biological knowledge-based approaches for the analysis of genome-wide expression profiles using gene sets and pathways,” Statistical Methods in Medical Research, vol. 18, pp. 577–593, Dec 2009.

[12] F. Maleki, K. Ovens, D. J. Hogan, and A. J. Kusalik, “Gene Set Analysis: Challenges, Opportunities, and Future Research,” Front Genet, vol. 11, p. 654, 2020.

[13] H. Yamamoto, T. Fujimori, H. Sato, G. Ishikawa, K. Kami, and Y. Ohashi, “Statistical hypothesis testing of factor loading in principal component analysis and its application to metabolite set enrichment analysis,” BMC Bioinformatics, vol. 15, p. 51, Feb 2014.

[14] M. V. DiLeo, G. D. Strahan, M. den Bakker, and O. A. Hoekenga, “Weighted correlation network analysis (WGCNA) applied to the tomato fruit metabolome,” PLoS One, vol. 6, no. 10, p. e26683, 2011.

[15] X. Zhan, A. D. Patterson, and D. Ghosh, “Kernel approaches for differential expression analysis of mass spectrometry-based metabolomics data,” BMC Bioinformatics, vol. 16, p. 77, Mar 2015.

[16] E. A. Franzosa, A. Sirota-Madi, J. Avila-Pacheco, N. Fornelos, H. J. Haiser, S. Reinker, T. Vatanen, A. B. Hall, H. Mallick, L. J. McIver, J. S. Sauk, R. G. Wilson, B. W. Stevens, J. M. Scott, K. Pierce, A. A. Deik, K. Bullock, F. Imhann, J. A. Porter, A. Zhernakova, J. Fu, R. K. Weersma, C. Wijmenga, C. B. Clish, H. Vlamakis, C. Huttenhower, and R. J. Xavier, “Gut microbiome structure and metabolic activity in inflammatory bowel disease,” Nat Microbiol, vol. 4, pp. 293–305, Feb 2019.

[17] Q. Feng, Z. Liu, S. Zhong, R. Li, H. Xia, Z. Jie, B. Wen, X. Chen, W. Yan, Y. Fan, Z. Guo, N. Meng, J. Chen, X. Yu, Z. Zhang, K. Kristiansen, J. Wang, X. Xu, K. He, and G. Li, “Integrated metabolomics and metagenomics analysis of plasma and urine identified microbial metabolites associated with coronary heart disease,” Sci Rep, vol. 6, p. 22525, Mar 2016.

[18] D. Jiang, C. R. Armour, C. Hu, M. Mei, C. Tian, T. J. Sharpton, and Y. Jiang, “Microbiome Multi-Omics Network Analysis: Statistical Considerations, Limitations, and Opportunities,” Front Genet, vol. 10, p. 995, 2019.

[19] J. Han, J. Meng, S. Chen, and C. Li, “Integrative analysis of the gut microbiota and metabolome in rats treated with rice straw biochar by 16S rRNA gene sequencing and LC/MS-based metabolomics,” Sci Rep, vol. 9, p. 17860, 11 2019.

[20] S. Kim, K. A. Sohn, and E. P. Xing, “A multivariate regression approach to association analysis of a quantitative trait network,” Bioinformatics, vol. 25, pp. i204–212, Jun 2009.

[21] S. Lee, M. J. Emond, M. J. Bamshad, K. C. Barnes, M. J. Rieder, D. A. Nickerson, D. C. Christiani, M. M. Wurfel, and X. Lin, “Optimal unified approach for rare-variant association testing with application to small-sample case-control whole-exome sequencing studies,” American Journal of Human Genetics, vol. 91, pp. 224–237, Aug 2012.

[22] J. A. Koziol and M. D. Perlman, “Combining independent chi-squared tests,” Journal of the American Statistical Association, vol. 73, no. 364, pp. 753–763, 1978.

[23] R. H. Berk and D. H. Jones, “Goodness-of-fit test statistics that dominate the kolmogorov statistics,” Zeitschrift für Wahrscheinlichkeitstheorie und verwandte Gebiete, vol. 47, no. 1, pp. 47–59, 1979.

[24] D. Donoho and J. Jin, “Higher criticism for detecting sparse heterogeneous mixtures,” The Annals of Statistics, vol. 32, no. 3, pp. 962–994, 2004.

[25] C. M. Carpenter, W. Zhang, L. Gillenwater, C. Severn, T. Ghosh, R. Bowler, K. Kechris, and D. Ghosh, “Pairkat: A pathway integrated regression-based kernel association test with applications to metabolomics and copd phenotypes,” PLOS Computational Biology, vol. 17, pp. 1–22, 10 2021.

[26] L. Dwyer-Lindgren, P. Kendrick, Y. O. Kelly, M. M. Baumann, K. Compton, B. F. Blacker, F. Daoud, Z. Li, F. Mouhanna, H. Nassereldine, C. Schmidt, D. O. Sylte, S. I. Hay, G. A. Mensah, A. M. poles, E. J. rez Stable, C. J. L. Murray, and A. H. Mokdad, “Cause-specific mortality by county, race, and ethnicity in the USA, 2000-19: a systematic analysis of health disparities,” Lancet, vol. 402, pp. 1065–1082, Sep 2023.

[27] S. Yusuf, S. Reddy, S. Ounpuu, and S. Anand, “Global burden of cardiovascular diseases: part I: general considerations, the epidemiologic transition, risk factors, and impact of urbanization,” Circulation, vol. 104, pp. 2746–2753, Nov 2001.

[28] M. L. Daviglus, A. Pirzada, and G. A. Talavera, “Cardiovascular Disease Risk Factors in the Hispanic/Latino Population: Lessons From the Hispanic Community Health Study/Study of Latinos (hchs/sol),” Progress in Cardiovascular Diseases, vol. 57, no. 3, pp. 230–236, 2014. Cardiovascular Diseases in Hispanics.

[29] S. Sharma and P. Tripathi, “Gut microbiome and type 2 diabetes: where we are and where to go?,” The Journal of Nutritional Biochemistry, vol. 63, pp. 101–108, 2019.

[30] A. Trikkalinou, A. K. Papazafiropoulou, and A. Melidonis, “Type 2 diabetes and quality of life,” World Journal of Diabetes, vol. 8, pp. 120–129, Apr 2017.

[31] J. N. Sampson, S. M. Boca, X. O. Shu, R. Z. Stolzenberg-Solomon, C. E. Matthews, A. W. Hsing, Y. T. Tan, B. T. Ji, W. H. Chow, Q. Cai, D. K. Liu, G. Yang, Y. B. Xiang, W. Zheng, R. Sinha, A. J. Cross, and S. C. Moore, “Metabolomics in epidemiology: sources of variability in metabolite measurements and implications,” Cancer Epidemiology, Biomarkers Prevention, vol. 22, pp. 631–640, Apr 2013.

